# Evolutionary Insights into Muscle Fiber Distribution in the Twin Tails of Ornamental Goldfish

**DOI:** 10.1101/2024.06.03.597082

**Authors:** Kinya G Ota, Gembu Abe, Chen-Yi Wang, Ing-Jia Li, Paul Gerald Layague Sanchez, Tzu-Chin Chi

## Abstract

Twin-tail ornamental goldfish have a bifurcated caudal fin with a morphology that is extremely diverged from the conventional body plan of the vertebrates. Here, we investigate the musculoskeletal histology of this bifurcated caudal fin. From some of the investigated twin-tail goldfish, we found a twin-tail goldfish specific muscle (hereafter referred to as the “medial caudal muscle”) between left and right bifurcated caudal fin skeletons. Our immunohistochemical analyses revealed that the medial caudal muscle showed laterally biased distribution patterns of the slow and fast muscle fibers. Similar distribution patterns were also commonly observed in several deep muscles of wild-type goldfish as well as zebrafish, suggesting that these muscle fiber distribution patterns are formed by the same molecular developmental mechanisms even though their morphologies are highly diverged. These findings provide empirical evidence to consider how the histological features of a newly emerged morphology are influenced by selective pressures and pre-existing developmental mechanisms.

## BACKGROUND

Ornamental twin-tail goldfish have a bifurcated caudal fin. This goldfish has been deliberately selected by breeders and fanciers since the Ming dynasty for its aesthete appeal (Chen, 1954, 1956; Ota, 2021; Ota and Abe, 2016). Owing to exceptional morphological characteristics (Abe et al., 2014; Li et al., 2019; Watase, 1887), this anatomical feature has been described in previous research. These studies revealed that the bifurcated caudal fin consists of the duplicated caudal skeletons and muscles (Fig. 1). Although several twin-tail mutants have been reported, such a morphology has not been genetically fixed in any other population besides twin-tail goldfish (see Korschelt, 1907; Tyler, 1970). The bifurcated caudal fin of twin-tail goldfish suggests that the basic architecture of caudal musculoskeletal systems underwent significant changes in the ornamental goldfish lineage throughout the domestication process (Abe et al., 2014; Flammang, 2014; Kardong, 2006; Lauder, 1989; Liem et al., 2001). That is to say, artificial selective pressures on the ornamental goldfish appear to have altered the evolutionarily highly conserved developmental mechanisms relating to musculoskeletal development (Abe et al., 2014).

**Fig 1.**
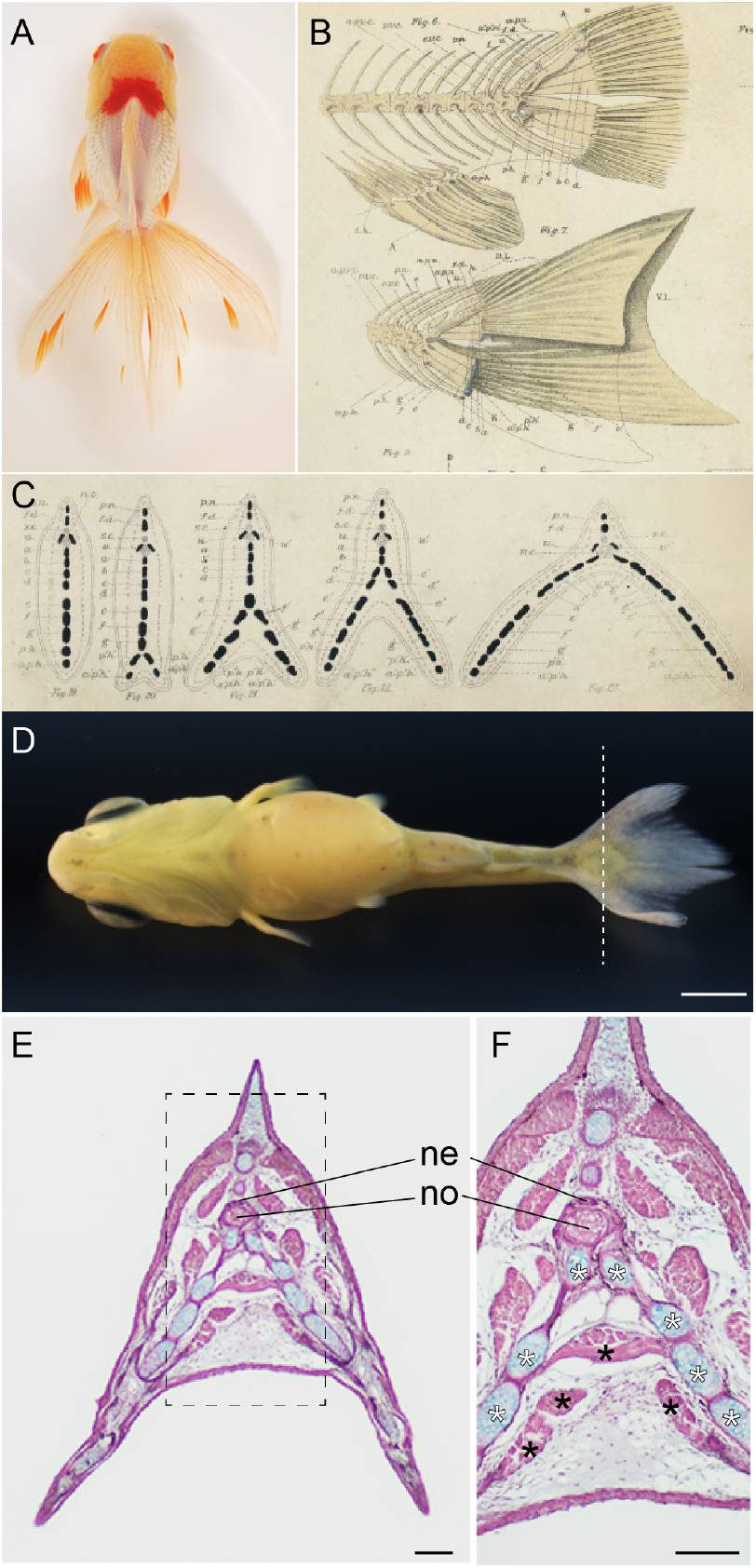
Bifurcated caudal fin and the medial caudal muscles. (A). Dorsal view of *Oranda* strain. (B, C). Drawing of the caudal region of common carp and twin-tail goldfish (B), and schematic view of the transverse section of goldfish caudal fin (C) by Watase (1887). (D). Ventral view of twin-tail goldfish larvae. (E). The transverse section of the caudal fin of twin-tail goldfish in panel D. Sectioned level is indicated by the dashed line on panel D. (F). The magnified view of panel E. The magnified region is indicated by the dashed line box in panel E. White asterisks indicate the caudal skeletons at the ventral side. The black asterisks indicate the medial caudal muscle. Abbreviations: ne, neural tube; no, notochord. Scale bars = 1 mm (D), 0.1mm (E, F). Panels A, D, E, F adapted from Li et al. (2019) under [Creative Commons Attribution 4.0 International License](https://creativecommons.org/licenses/by/4.0/).

The twin-tail phenotype is retained in the ornamental goldfish population mainly due to its attractiveness for breeders and fanciers. It is known that the bifurcated caudal musculoskeletal system in the twin-tail goldfish stems from the highly modified dorsal-ventral patterning deriving from the homozygous locus of a stop codon mutation allele in the duplicated *chordin* gene, referred to as the *chdS*^*E127X*^ allele (or *chdA*^*E127X*^ in the previous report) (Abe et al., 2014; Kon et al., 2020). This mutation alters the gene’s original function, resulting in ventralized early embryos, the formation of bifurcated caudal fin folds at the embryonic stage, and ultimately, the emergence and stable fixation of various ornamental twin-tail goldfish strains (Abe et al., 2014; Chen et al., 2022).

Anatomical changes in the emergence of twin-tail goldfish also imply that the *chordin* gene mutation influence not only early development and patterning but also histogenesis. Specifically, we identified muscle that had not been reported in other teleost species (Li et al., 2019); in this study, we refer to this unique muscle as the “medial caudal muscle” (Fig. 1DEF). Positioned between the bifurcated left and right caudal skeletons, these twin-tail goldfish-specific muscles raise several questions. For instance, it prompts inquiry into which muscle in the wild-type goldfish is most closely related to the medial caudal muscle. This query can be reframed as follows; is the medial caudal muscle entirely novel and unique within the lineage of twin-tail goldfish, or are there comparable features between the medial caudal muscles and the conventional muscles in wild-type goldfish? Although the medial caudal muscle exhibits morphological uniqueness in twin-tail goldfish, could a comparable muscle be identified in wild-type goldfish by examination at a finer resolution, such as at the level of muscle tissue characteristics?

Muscle tissues of vertebrate species are heterogeneous in their consisting of muscle fibers in general. Although there are several different ways to classify and identify the muscle fiber types of the vertebrate species (see Luna et al., 2015; Keenan and Curie, 2019), the teleost muscle fibers are categorized into fast and slow muscle fibers (Bernal et al., 2001; Devoto et al., 1996; Du et al., 1997; Fierstine and Walters, 1968; Shadwick et al., 2002; Stickney et al., 2000). These muscle fibers show well-arranged distribution patterns at the mid-trunk region (Devoto et al., 1996; Kardong, 2006; Liem et al., 2001; Nakae et al., 2014; Westneat and Wainwright, 2001). Fast muscle fibers are located on the medial side, and on the other hand, the slow muscle fibers are distributed on the bilateral surface at the mid-trunk region in the majority of the teleost species. While the distribution patterns of the fast and slow muscle fibers are exceptionally highly modified in the lineage of the tuna fish group due to their swimming behavior, it is generally recognized that the same distribution patterns of these muscle fibers are highly conserved among major teleost species (Bernal et al., 2001; Fierstine and Walters, 1968; Shadwick et al., 2002).

The conserved distribution patterns of the different types of muscle fibers have also been investigated at the levels of developmental biology. Particularly, the differentiation and migration patterns of the slow muscle fibers have been intensively studied in zebrafish at the molecular level (Devoto et al., 1996; Du et al., 1997; Keenan and Currie, 2019; Stickney et al., 2000). These studies revealed that slow and fast muscle fibers differentiate from embryonic somite cells. Initially, slow muscle primordial cells are distributed to adaxial regions of segmentally arranged somites, and these cells migrate to the lateral side and form the slow muscle fibers on the lateral surface at the trunk region (Devoto et al., 1996; Du et al., 1997; Keenan and Currie, 2019; Stickney et al., 2000). Notably, critical signals from the notochord, including hedgehog signaling, play a pivotal role in these differentiation processes. These suggest that modifications to the early developmental process may impact the differentiation of these muscle fibers in the bifurcated caudal fin of the twin-tail goldfish.

Presumably due to the complexity of the muscle arrangement and structure in the caudal region, there has been limited immunohistochemical research conducted on muscle fibers in this area. In fact, while early research showed that the caudal muscle of teleost species contains both slow and fast muscles, their distribution patterns in muscle tissues and their relative location with the other skeletal tissues have not been well investigated yet (Flammang and Lauder, 2008, 2009; Kryvi et al., 2021; Nag, 1972). Thus, in this study, we conducted a detailed immunohistochemical analysis of the musculoskeletal system in the twin-tail goldfish, especially, focusing on the caudal region. Aiming to examine how histological features were changed/conserved in the twin-tail goldfish, we performed immunohistochemistry focusing on the muscle fiber distribution patterns. We here carefully examined the distribution patterns of the slow and fast muscle fibers in the twin-tail goldfish and compared them with those in the wild-type goldfish and those in zebrafish.

Moreover, we examined two ornamental domesticated goldfish (including *Ryukin* and *Oranda* goldfish strain), and the lab strain goldfish derived from *chdS* mutant parents. We also deduce how histological features of the musculoskeletal system could be modified by drastic morphological changes within a short period due to genetic mutations and artificial selection, more generally aiming to provide implications for understanding large-scale morphological evolution and the emergence of novel phenotypes.

## MATERIALS AND METHODS

### Goldfish and zebrafish

All the goldfish strains were maintained in Yilan Marine Research Station. The wild-type goldfish strain was derived from the single-tail common goldfish of the Taiwanese and Japanese populations. The single-tail goldfish strain parents were genotyped at the *chdS* locus (*chdA* in Abe et al., 2014) and *chdS*^*wt/wt*^, *chdS*^*wt/E127X*^, and *chdS*^*E127X/E127X*^ individuals were separately maintained in our aquarium facility. To reproduce the wild-type goldfish, we conducted artificial fertilization of *chdS*^*wt/wt*^ genotype males and females. Goldfish showing *chordin* gene mutated phenotypes were reproduced by crossing *chdS*^*E127X/E127X*^ and/or *chdS*^*wt/E127X*^ parents. From the segregant population containing different *chdS* genotype, the progenies showing the bifurcated caudal fin fold were collected at 3-4 days post-fertilization (dpf). The *Ryukin* and *Oranda* strain progenies were obtained by crossing their parents purchased from local ornamental fish distributors based in Taiwan. These progenies were obtained by artificial fertilization as described previously (Li et al., 2019). Zebrafish adult specimens were derived from our lab strain. The wild-type zebrafish individuals were derived from the lab strain originally established from the progenies of the AB strains provided by Zebrafish Core Facility at Academia Sinica, Taiwan (TZCAS). All experiments involving these fish have been approved by the Institutional Animal Care and Use Committee (IACUC) at Academia Sinica (Protocol ID: #19-11-1351 #20-06-1480 #22-11-1922 #23-10-2073).

### Maintenance of larvae and juveniles

Early stage larvae were moved from plastic dishes (9 cm) to plastic tanks (3000 ml). The plastic tanks are located in the aquarium system with an overflow system. The quality of water in the aquarium system was automatically adjusted to 200 to 300 *µ*S/cm in conductivity, pH 6.5 to 7.5, and 24°C to 26°C. Progenies were fed with live food (paramecium and/or brine shrimp) and dry food at least once per day; the type of feed depended on the size of the progenies. Prot stage larvae were fed with paramecium. After Prot stage, larvae were mainly fed with brine shrimp at least once per day and supplemented with paramecium, algae, and dry food to minimize the risk of starvation and nutritional deficiency (Li et al., 2015, 2019; Tsai et al., 2013).

### Histological analyses

Fish specimens were anesthetized with MS-222 (E10521, Sigma) and fixed using Bouin’s solution (HT10132, Sigma). The fixed specimens were soaked in 70% ethanol and photographed under the stereomicroscope (SZX16 with DP80, Olympus). After dehydration, specimens were cleared in Lemosol A (5989-27-5, FUJIFILM Wako Pure Chemical Corporation) embedded in paraffin, and sectioned to 5 *µ*m using a microtome (RM2245, Leica). The sections were placed on microscope slides (Platinum-pro, Matsunami). For the conventional histological analyses, the section was deparaffinized with Histo-Clear II (HS-202, National Diagonistics), gradually hydrated with ethanol series, and stained as described previously (Li et al., 2019). For immunohistochemistry, the section was incubated at 65°C for 20 min in a paraffin oven, deparaffinized with Histo-Clear II, and immersed in 100% ethanol. The deparaffinized slide was immersed in 1% hydrogen peroxide/methanol solution for more than 1 hour, and hydrated with ethanol series. The hydrated slide is blocked in 3% skim milk PBST for 1 hour and incubated with primary antibody (1:100) diluted in Superblock Blocking Buffer in TBS (#37535, Thermo Fisher Scientific) at room temperature overnight. The primary antibodies used are F59 (sc-32732, Santa Cruz Biotechnology Inc.) and F310 (Developmental Studies Hybridoma Bank). On the following day, the slides were washed with PBST five times for 3 min each and incubated with the secondary antibody (Goat Anti-Mouse IgG H&L [ab205719, Abcam]) (1:1000) at room temperature for more than six hours. After the secondary antibody incubation, the slides were washed five times each in PBST for 3 min each. The signals were detected by Thermo Scientific Pierce Metal Enhanced DAB substrate kit (#34065, Thermo Scientific).

The counterstaining was performed using Alcian blue, hematoxylin, and Giemsa stains. The staining conditions were as follows: microscope slides were immersed in 0.1% Alcian blue (A5268, Sigma) aqueous solution for 30 seconds to 1 minute, then briefly rinsed with water to remove excess stain before being differentiated with 0.01% HCL in 70% ethanol. Next, they were stained with hematoxylin solution (MHS32, Sigma) for 30 seconds to 1 minute, followed by staining with 5% Giemsa solution (Sigma, GS500) in PBS for 15 minutes. The slides were then rinsed with running water for about 5 minutes, differentiated in 80% ethanol for approximately 5 seconds, and finally washed with running water for an additional 30 minutes. Hematoxylin and Alcian blue staining allow for the differentiation of cell nuclei and cartilage tissue. Additionally, Giemsa staining strongly colors melanocytes in a bluish-black hue, making it possible to distinguish them from the brown coloration of DAB staining (Luiza Silveira et al., 2020). The slide was dehydrated with ethanol series, immersed in Lemosol A, and sealed by coverslip with Entellan new mounting media (107961, Sigma-Aldrich). The slides were observed under the microscope (BX43 with DP27, Olympus). Identification and nomenclature of skeletons and muscles were based on Kardong (2006); Lauder (1989); Liem et al. (2001); Siomava and Diogo (2018); Winterbottom (1973).

## RESULTS

### The wild-type goldfish

To investigate the basic characteristics of slow and fast muscle fibers in the wild-type goldfish caudal muscles, we first conducted immunohistochemistry by using two distinct antibodies—F310 (the fast muscle fiber antibody) and F59 (the slow muscle fiber antibody) (Fig. 2, 3, and 4). The histological sections from the anterior caudal region (or the caudal peduncle) of the pelvic fin ray stage larva showed the distribution patterns of the muscle fibers of a typical teleost species (Fig. 2A-D2). The predominant portion of the muscle located on the medial side comprised fast muscle fibers, with its bilateral sides enveloped by slow muscle fibers (Fig. 2C-2D). The boundary between these muscle fibers was discernible, indicating the suitability of F310 and F59 antibodies for the immunohistochemical analysis in goldfish.

**Fig 2.**
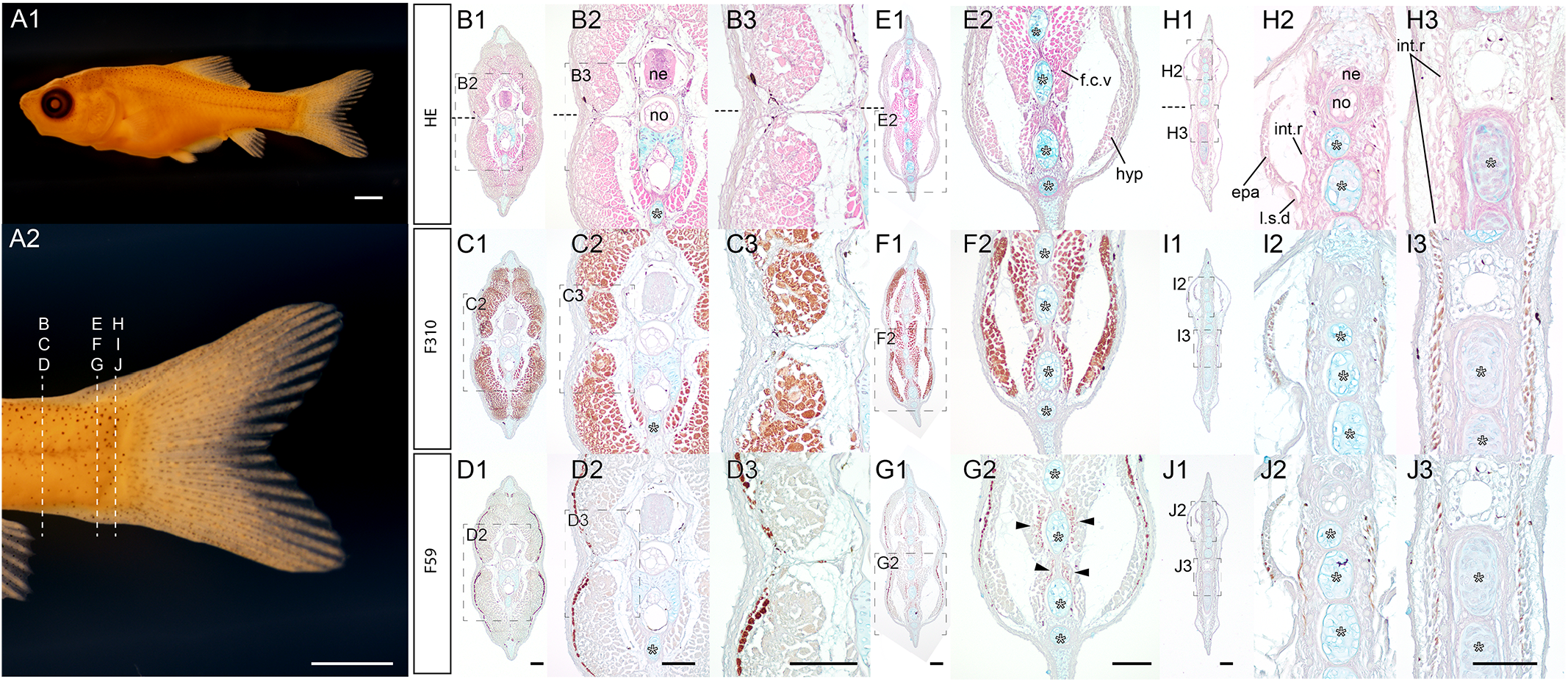
Transverse section of a wild-type goldfish larva at the pelvic fin ray stage. (A). Lateral view of goldfish fixed with Bouin’s fixative (#2021-0517-09-22dpf-B01ZWJZWJ; 22 dpf and 9.95mm in standard length). (A). Whole lateral view (A1) and magnified caudal view (A2). (B-J). Transverse sections at the anterior level (B-D), the mid-level (E-G), and the most posterior level (H-J) of the caudal region. The same sections are indicated by the same Roman letters and the magnified views are identified by the plural numeric suffix on the left upper corner of the panels. Dashed boxes are indicated the magnified regions. The upper, middle, and lower panels of the histological sections are conventional histological sections, immunohistochemistry with F310 antibody (the fast muscle fiber), and with F59 antibody (the slow muscle fiber), respectively. White asterisks indicate the ventral caudal skeleton complex including, hypural, parhypural, and hermal spines. Approximate sectioned levels are indicated by dashed lines in panel (A2). Horizontal myoseptum is indicated by black dashed lines in panels B1, B2, and B3. The slow muscle fibers are indicated by black arrowheads in panel G2. The conventional histology, F310, and F59 antibodies-stained sections in the same column are derived from the adjacent sections. Abbreviations: epa, epaxial muscle; f.c.v, flexor caudalis ventralis; hyp, hypaxial muscle; int.r, interradials; l.s.d, lateralis superficialis dorsalis; ne, neural tube; no, notochord. Scale bars = 1mm (A1, A2), 100*µ*m (D1, D2, D3, G1, G1, G2, J1, J3). Histological sections in the same column have the same magnification.

**Fig 3.**
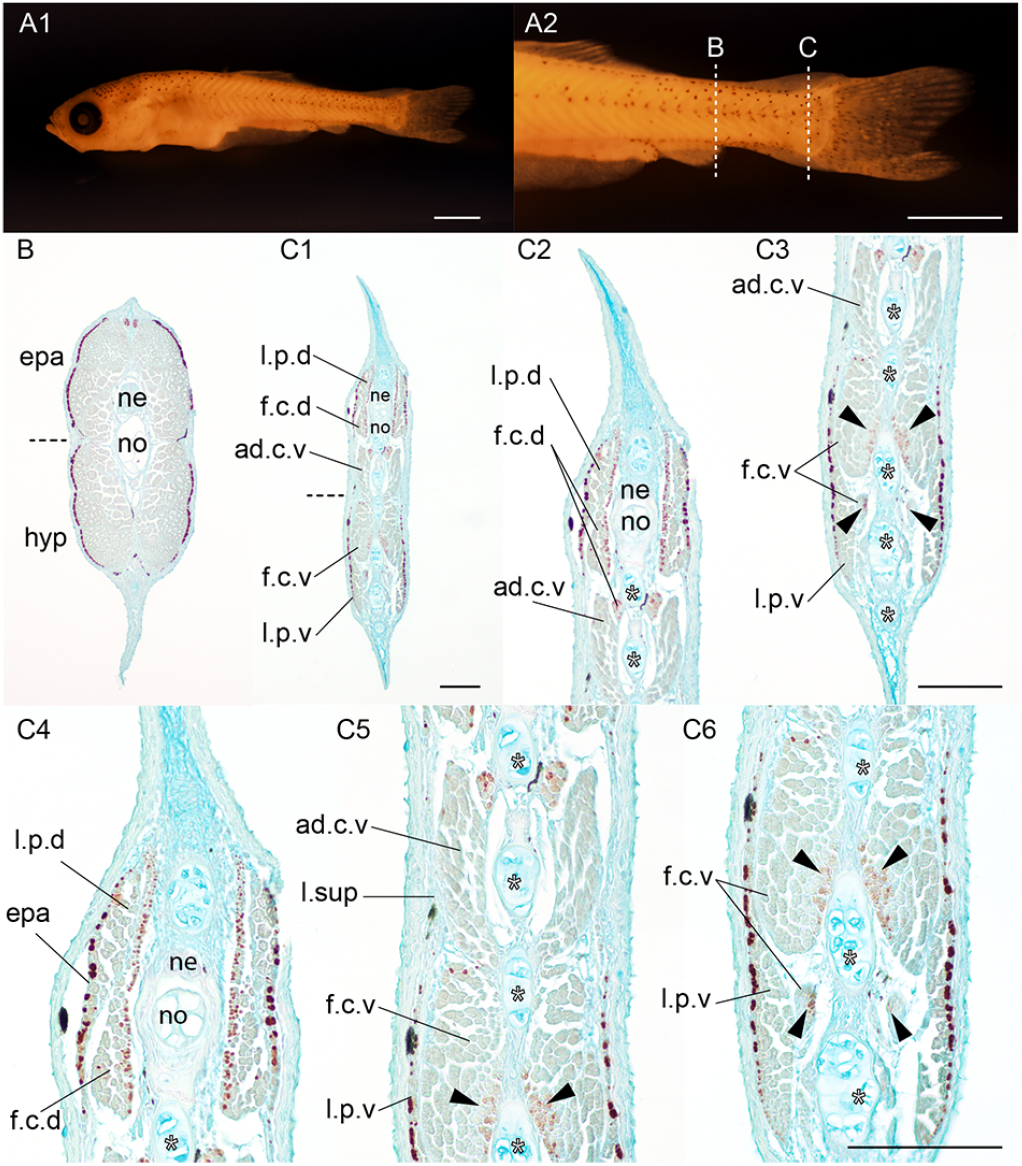
Transverse section of a wild-type goldfish larva at the late pelvic fin bud stage. (A). Whole lateral view (A1) and magnified view of caudal level (A2) of the goldfish larvae (#2020-0406-01-Bzwj, 7.97 mm, 26 dpf). (B, C). Transverse sections immunostained with the slow muscle fibers specific antibody (F59). (C2, C3). Medium magnification views of dorsal (C2) and ventral (C3) sides at the caudal level sections. (C4-6). High magnification views of dorsal (C4), mid (C5), and ventral regions at the caudal level sections. Sectioned levels are indicated by dashed lines in panel A2. The horizontal myoseptum is indicated by black dashed lines in panels B and C1. White asterisks indicate the ventral caudal skeleton complex including pural, hypural, and hermal spines. Black arrowheads indicate slow muscle fibers in the flexor caudalis ventralis. Abbreviations: ad.c.v, adductor caudalis ventralis; epa, epaxial muscle; f.c.d, flexor caudalis dorsalis; f.c.v, flexor caudalis ventralis; hyp, hypaxial muscle; l.p.d, lateralis profundus dorsalis; l.p.v, lateralis profundus ventralis; l.sup, lateralis superficials; ne, neural tube; no, notochord. Scale bars = 1mm (A1, A2), 100*µ*m (C1, C3, C6). Panel B and C1, C2 and C3, and panels of the third row have the same magnifications.

**Fig 4.**
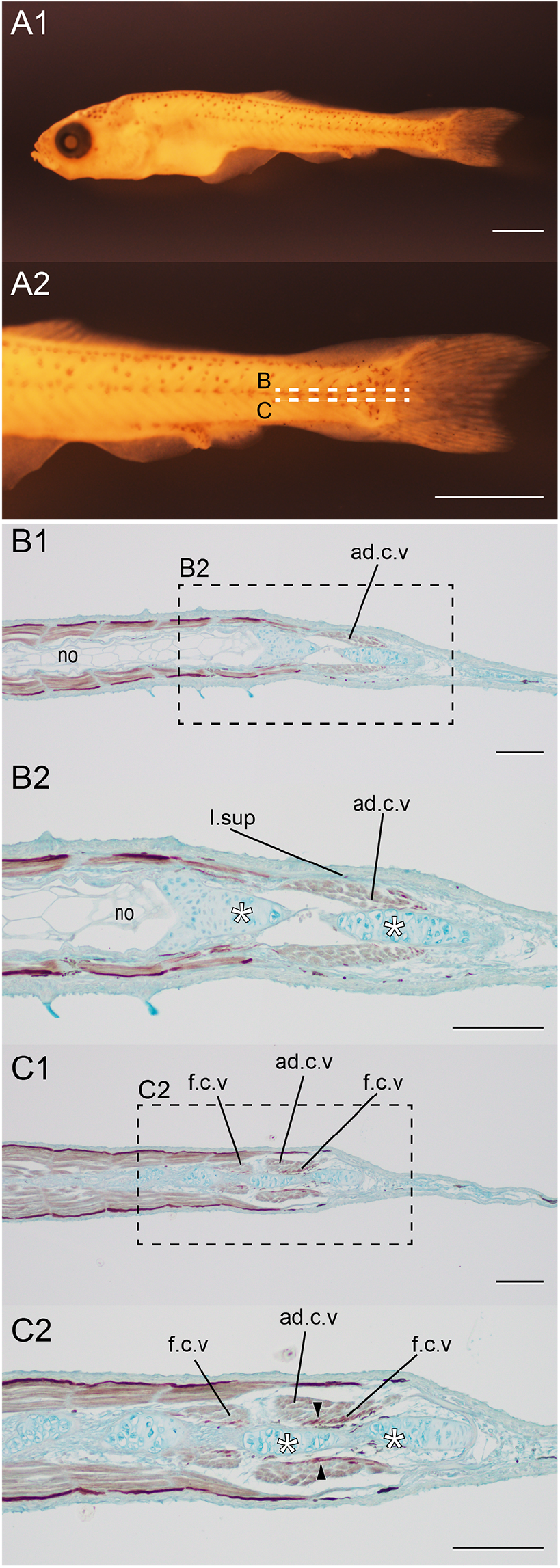
Horizontal section of the caudal region of a wild-type goldfish larva at pelvic fin bud stage. (A). The whole (A1) and the magnified (A2) view of the goldfish (#2020-0406-01C; SL=8.07 mm; 26 dpf). (B, C). Sections immunostained with the slow muscle fiber specific antibody (F59) in the caudal region. The sectioned levels are indicated in panel A2. The magnified views of B1 and C1 are shown in B2 and C2, respectively. White asterisks in B2 and C2 indicate axial skeleton. Slow muscle fibers of flexor caudals ventralis are indicated by black arrowheads. Abbreviations: ad.c.v, Adductor caudalis ventralis; f.c.v, flexor caudalis ventralis; l.sup, lateralis superficials. Scale bars = 1mm (A), 100 *µ*m (B, C).

At the more posterior caudal levels, the distribution pattern of these muscle fibers becomes more intricate (Fig. 2E-2G). While fast muscle fibers on the superior layers exhibit similar distribution patterns to those observed at the caudal peduncle levels, the deeper layers of muscle fibers (flexor caudalis ventralis, f.c.v) contain slow muscle fibers (Fig. 2E-2G). Signal corresponding to slow muscle fibers is observed from the most medial side of the section, where the muscle fibers attaching to the caudal axial skeleton are detected (the black arrowheads of Fig. 2G2). Towards more posterior regions, the epaxial (epa) muscles and interradial (int.r) muscles display two types of muscle fibers (Fig. 2H-2J). We could observe that the fast and slow muscle fibers distribute in the surface and deeper sides, respectively (Fig. 2IJ). Using adjacent sections, we confirmed that muscle fibers showing immunoreactivity with both F59 and F310 antibodies were not detected, and no muscle displaying a random mixture of slow and fast fibers was observed. In other words, these results demonstrate that the antibodies used to stain different types of muscle fibers without problematic cross-immunoreactivity, and even when using the F59 antibody alone, they provide insight into the distribution of slow and fast muscle fibers.

For a more detailed examination of slow muscle fiber distribution patterns, we conducted immunohistochemistry with F59 antibody in a serial section of the wild-type goldfish larva (Fig. 3). Strong signals from the F59 antibody revealed the presence of slow muscle fibers on the superior parts at both the peduncle and posterior caudal levels, demonstrating the consistency of our immunohistochemistry (Fig. 3A2, 3B, 3C1). Notably, F59 antibody signals were observed in the deeper parts of muscles on both dorsal and ventral sides at the posterior level of the caudal region. These histological images enabled the identification of slow muscle fiber locations in various parts of the caudal muscles (Fig. 3C1-3C6). The bilateral sides of the surface musculatures—although their boundaries are uncertain, likely involving lateralis profundus dorsalis/ventralis (l.p.d/l.p.v), and lateralis superficialis dorsalis/ventralis (l.sup) —contained slow muscle fibers (Fig. 3C2-3C6). Conversely, in deep musculatures including flexor caudalis dorsalis (f.c.d) and flexor caudalis ventralis (f.c.v), slow muscle fibers were distributed on the medial side (Fig. 3C2-3C6).

Similar distribution patterns of slow muscle fibers were identified in horizontal sections of pelvic fin bud stage larvae (Fig. 4A). Slow muscle fibers were observed on the bilateral surface sides at the caudal peduncle level (Fig. 4B1, 4B2). Additionally, flexor caudalis ventralis (f.c.v) displayed relatively strong signals of F59 antibody toward the midline, indicating that the slow muscle fibers are positioned near the caudal skeleton (Fig. 4C1, 4C2). In total, several muscles of the caudal region differ significantly from that of the midtrunk region in the distribution patterns of slow and fast muscle fibers in the wild-type goldfish.

### Comparison with zebrafish

To examine whether the observed slow muscle distribution patterns are goldfish-specific characteristics or not, we further examined the slow muscle fiber distribution patterns in zebrafish adults (Fig. 5A). Our immunohistochemistry analysis indicated that the slow muscle fibers were also distributed at the medial regions of the deep muscles at the caudal level; the same results were observed in two different individuals (Fig. 5BC). The signals of the slow muscle antibody are detected at the bilateral side of the muscle fibers in the trunk muscles, and more significantly, in flexor caudalis ventralis (f.c.v), consistent with the results in wild-type goldfish (the black arrowheads in Fig. 2G2, 3C2, 3C4, 3C5, 3C6, 4C2, 5B2, 5B3, 5C2, and 5C3). As observed in the single-tail common goldfish, the muscle fibers proximate to the caudal axial skeletons in the flexor caudalis ventralis (f.c.v) tended to show the F59 antibody-positive muscle fibers (the black arrowheads in Fig. 5B2, 5B3, 5C2, and 5C3). Namely, different from those of the mid-trunk region, the distribution patterns of slow muscle fibers of the observed deep muscles in the caudal region are medially axially biased in zebrafish as well. This suggests that the bilaterally biased distribution patterns of the slow muscle fibers in the deep muscle at the caudal level are commonly conserved histological characteristics in these two teleost species.

**Fig 5.**
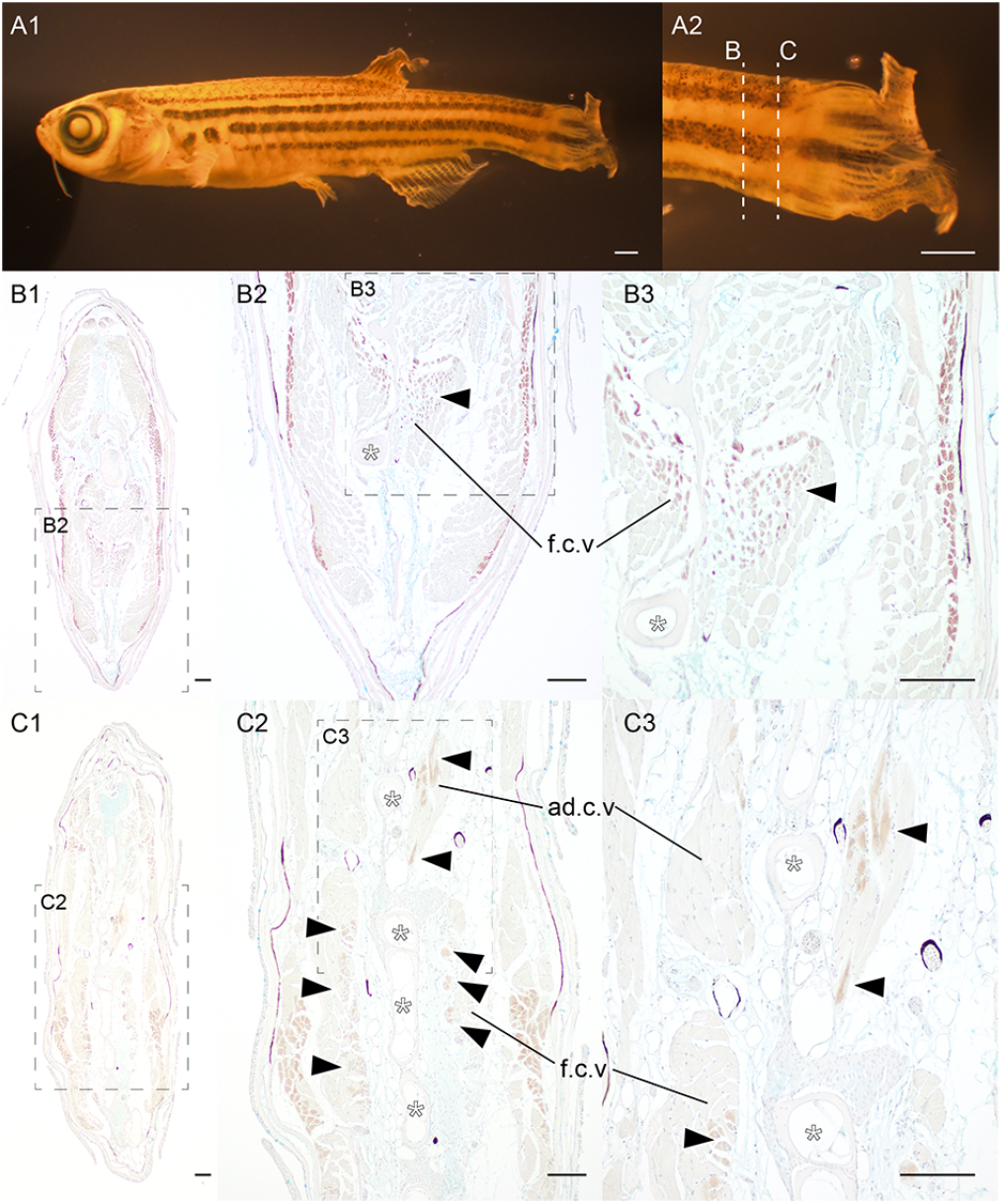
Slow muscle tissues at the caudal region of zebrafish. (A). The lateral view of zebrafish (2022-0607-ZF-labstrain, 582 dpf and 22.0 mm in standard length). (B, C). Transverse sections of immunostained with the slow muscle fiber specific antibody (F59) at caudal fin level. The magnified views of the regions are outlined by the dashed box with panel labels. Panels B and C are derived from two different individuals. Approximate levels of the histological sections are indicated by dashed lines in panel A2. White asterisks indicate the ventral caudal skeleton complex. Black arrowheads indicate slow muscle fibers in flexor caudalis ventralis and adductor caudalis ventralis. Abbreviations: ad.c.v, adductor caudalis ventralis; f.c.v, flexor caudalis ventralis. Scale bars = 1mm (A), 100 *µ*m (B1-C3).

### Twin-tail ornamental goldfish

To find similar/different points between the wild-type and twin-tail goldfish, we conducted conventional histological analysis and immunohistochemistry in the pelvic fin ray stage *Ryukin* strain larva (Fig. 6A). Similar to the wildtype goldfish, histological analyses at the peduncle level of *Ryukin* goldfish showed robust signals from F310-positive fast muscle fibers throughout the major portion of muscular tissues, with the superior muscle layer instead exhibiting F59-positive slow muscle fibers (Fig. 6B-D). At the posterior level of the caudal region on the dorsal side, the muscles of *Ryukin* goldfish displayed several muscles reminiscent of those observed in the wild-type goldfish; lateralis profundus dorsalis (l.p.d), flexor caudalis dorsals (f.c.d), and adductor caudalis (ad.c.v) can be easily recognized (Fig. 6E-G).

**Fig 6.**
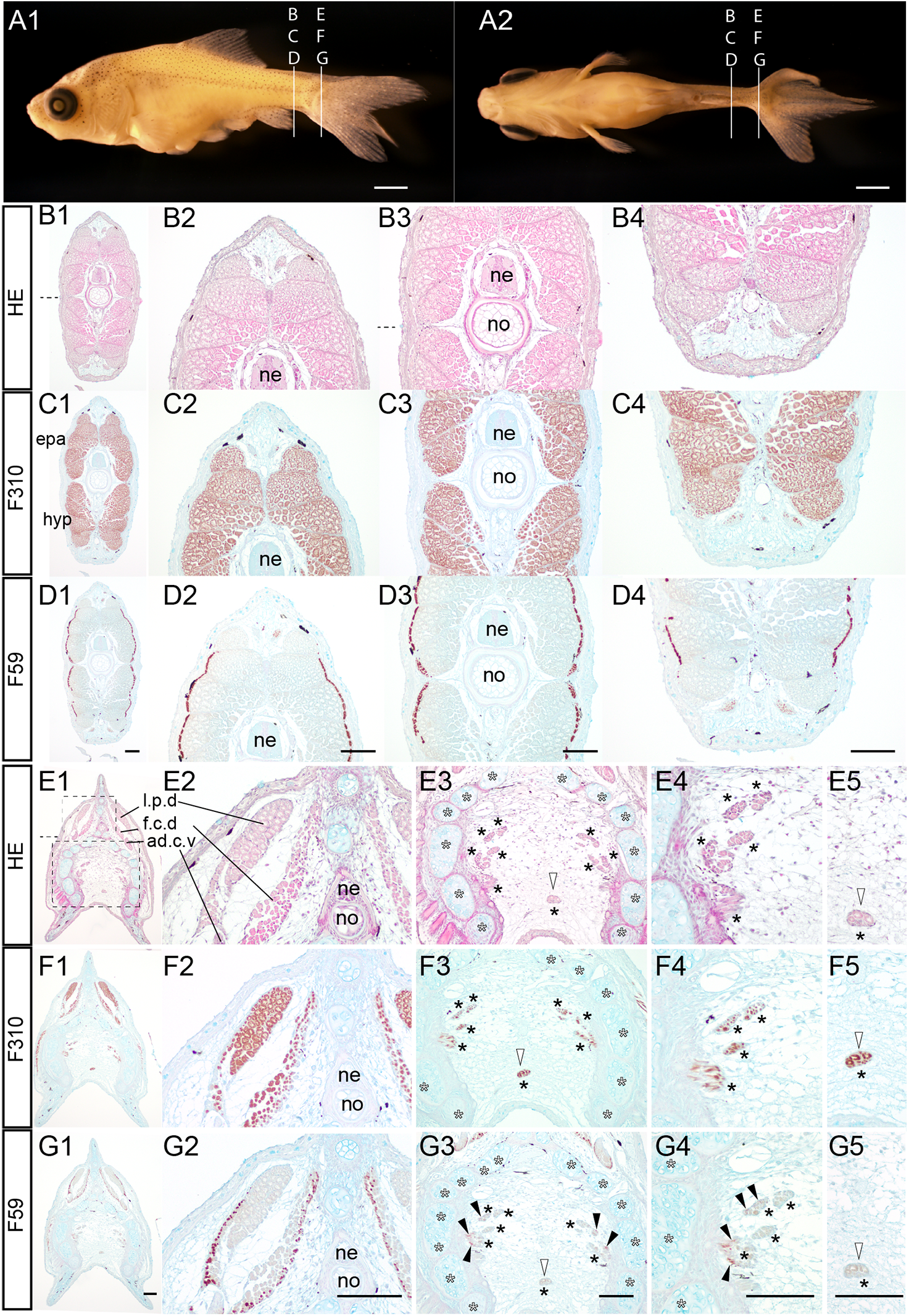
Transverse view of the trunk region of a twin-tail goldfish larva at the pelvic fin ray stage. (A). The lateral view (A1) and ventral view (A2) of *Ryukin* goldfish sample (#2022-0502-21-26dpdf-RY@02-0802-1A-6C-2A-#01; 8.84 mm; 26 dpf). (B-G). Transverse sections hematoxylin-eosin stained and immunostained with the fast muscle fiber specific antibody (F310), and the slow muscle fiber specific antibody (F59) at post-anal fin level (B-D) and caudal fin level (E-G). Panels in the second to fourth columns at post-anal fin levels show magnified views of the dorsal (B2, C2, D2), mid (B3, C2, D3), and ventral (B4, C4, D4) regions. Panels in the second and third columns at caudal levels showed magnified views of the dorsal (E2, F2, G2), and ventral (E3, F3, G3) regions. Magnified areas are outlined by dashed boxes in panel E1. Panels in the fourth and fifth columns at caudal levels showed high-magnified views of the left-ventral (E4, F4, G4) and mid-ventral (E5, F5, G5) regions. Horizontal myoseptum is identified by dashed lines in panels B1, B3 and E1. The muscular tissues located between the bifurcated caudal fin skeleton are indicated by black asterisks. White asterisks indicate the ventral caudal skeleton complex including pural, hypural, and hermal spines. Black arrowheads indicate slow muscle fibers in the flexor caudalis ventralis. Abbreviations: ad.c.v, adductor caudalis ventralis; epa, epaxial muscle; f.c.d, flexor caudalis dorsalis; f.c.v, flexor caudalis ventralis; hyp, hypaxial muscle; l.p.d, lateralis profundus dorsalis; ne, neural tube; no, notochord. Scale bars = 1 mm (A2), 100 *µ*m (D1, D2, D3, D4, G1 G2, G3, G4, G5). Adjacent histological sections in the same column have the same magnification.

In addition, on the ventral side, medial caudal muscle tissues were evident (black and white arrowheads in Fig. 6E-G), located in the intermediate regions of the bifurcated caudal skeletons (Fig. 6E-G), as previously reported (Li et al., 2019). While most of these muscle fibers are found close to the bifurcated caudal skeletons (the black asterisks in Fig. 6E3), a single muscle population, indicated by the white arrowhead in Fig. 6E3, seems to be isolated from the others and located on the middle sagittal plane of the body (Fig. 6E5, 6F5). The distribution patterns of the slow and the fast muscle fibers suggest that these twin-tail goldfish-specific muscular tissues contain both slow and fast muscle fibers (Fig. 6F4, 6F5, 6G4, 6G5). The slow muscle fibers in the medial caudal muscle are distributed near the caudal skeleton (the black arrowheads in Fig. 6G3, 6G4). Moreover, the same distribution patterns of the slow muscle fibers were observed in the horizontal plane of *Ryukin* strain larva (Fig. 7).

**Fig 7.**
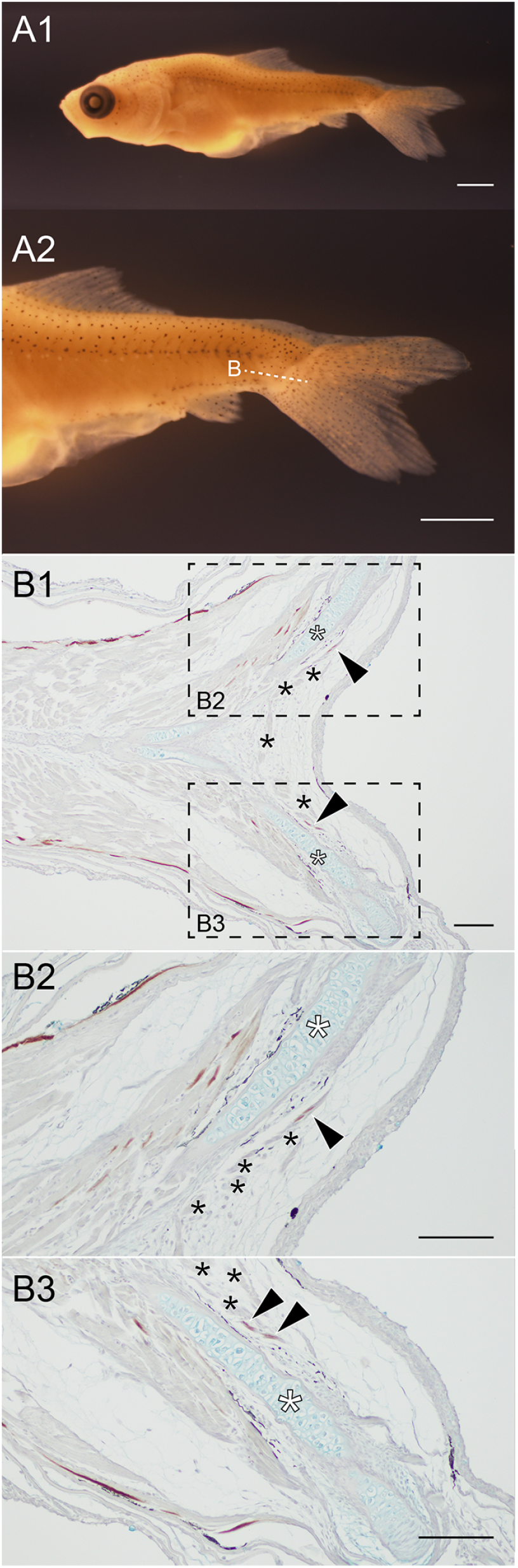
Horizontal view of the distribution of the slow muscle fibers at the caudal region of *Ryukin* larva. (A). Whole body (A1) and the magnified caudal region (A2) of *Ryukin* larva (2022-0502-21-26dof-@04, 26 dpf, 9.11 mm in standard length). (B). The wide (B1) and magnified views (B2, B3) of horizontal section stained with the slow muscle fiber specific antibody (F59). Magnified areas are indicated by dashed boxes in B1. The muscular tissues located between the bifurcated caudal fin skeleton are indicated by black asterisks. White asterisks indicate the ventral caudal skeleton complex including pural, hypural, and hermal spines. Black arrowheads indicate slow muscle fibers in the flexor caudalis ventralis. Sectioned levels are indicated on panel of A2. Scale bars = 1 mm (A1, A2), 100 *µ*m (B1, B2, B3).

To examine whether the same distribution patterns of the slow muscle fibers could be observed in the differnt types of twin-tail ornamental goldfish strain, we conducted the imunohistological analyses in the *Oranda* strain (Fig. 8A). Similar to the *Ryukin* strain (Fig. 6, 7), the strong signals were obtained in the lateralis profundus dorsalis of the *Oranda* strain (8B1-B2). More significantly, medial caudal muscles of this *Oranda* individual showed the slow muscle fibers (Fig. 8B3-B5). The signals are subtle in comparison with the investigated *Ryukin* individuals (Fig. 8B3-B5). However, the mediolaterally biased distribution patterns were observed in the medial caudal muscle, showing consistent results from *Ryukin* strain (Fig. 7).

**Fig 8.**
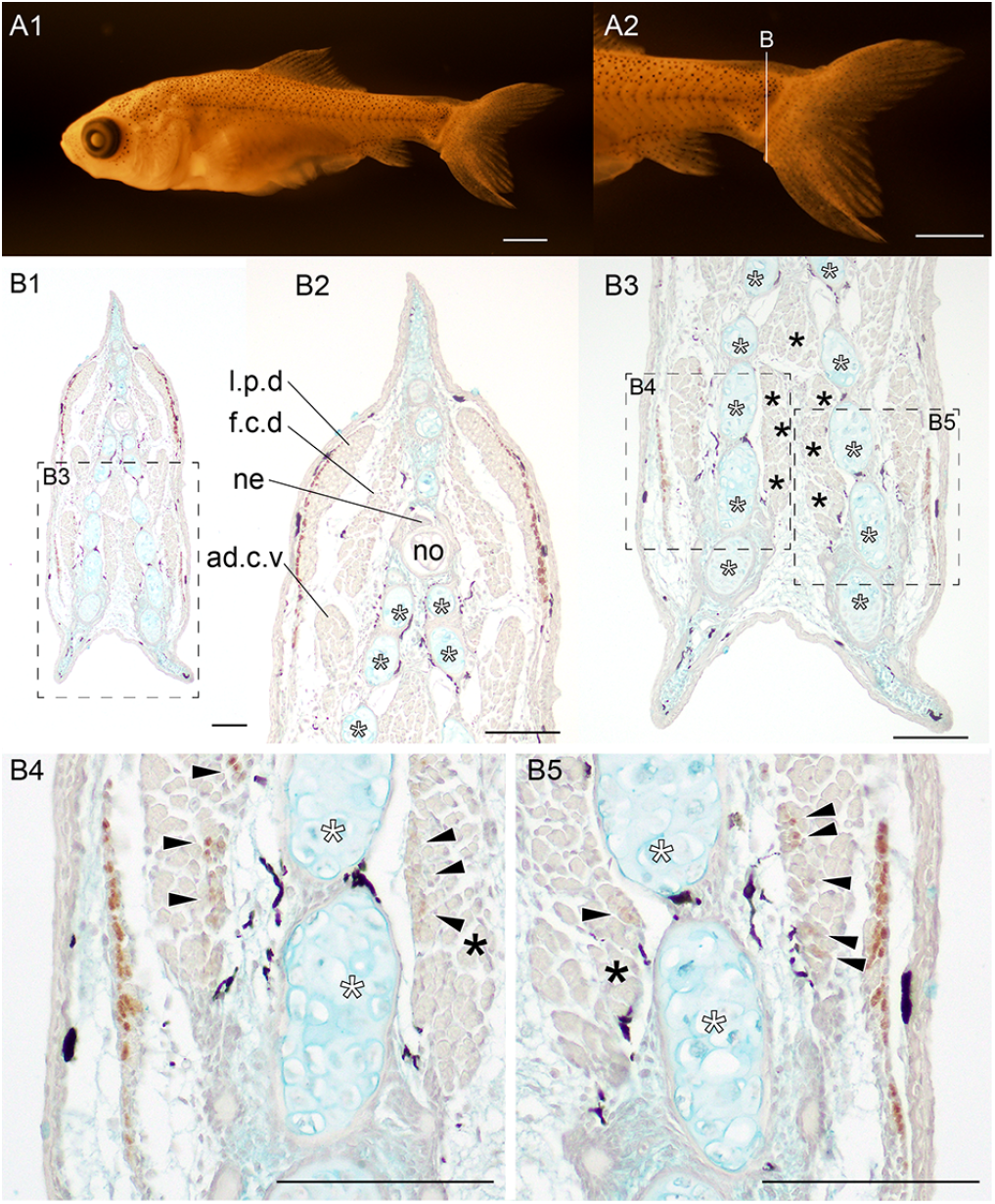
Distribution pattern of the slow muscle fibers in the *Oranda* strain goldfish. (A). Whole body (A1), and magnified lateral (A2) of goldfish (#2024- 0319-01-#03; 7.93 mm in the standard length, 31 dpf). The section at the caudal region was immunostained by the slow muscle fiber specific antibody (F59). (B). The sectioned levels are indicated in A2. White asterisks indicate the ventral caudal skeleton complex including pural, hypural, and hermal spines. Black asterisks indicate the medial caudal muscle. Abbreviations: ad.c.v, adductor caudalis ventralis; f.c.d, flexor caudalis dorsalis; l.p.d, lateralis profundus dorsalis; no, notochord; ne, neural tube. Scale bars = 1 mm (A1, A2), 100 *µ*m (B).

### *chdS* mutant lab strain progenies

We further examined the distribution patterns of slow and fast muscles in goldfish displaying various phenotypes of the caudal fin morphotype. Through artificial fertilization of the *chdS* mutant lab strain male and female, we successfully obtained seven juveniles exhibiting distinct morphologies in the caudal fin (MATERIALS AND METHODS) (Fig. 9-14). Based on the year of artificial fertilization, characteristic phenotype, and experimental ID, we designate them as follows: the “*23 single #03*” (#2023-0417-09-#03: Fig. 9), “*23 narrow twin #08*” (#2023-0417-09-#08: Fig. 10), “*23 wide twin #01*” (#2023-0417-09-#01: Fig. 11)”, “*23 twisted #04*” (#2023-0417-09-#04: Fig. 12), “*24 twin #01*” (#2024-0419- 03-#01: Fig. 13), and “*24 twin #02*” (#2024-0419-03-#02: Fig. 14).

**Fig 9.**
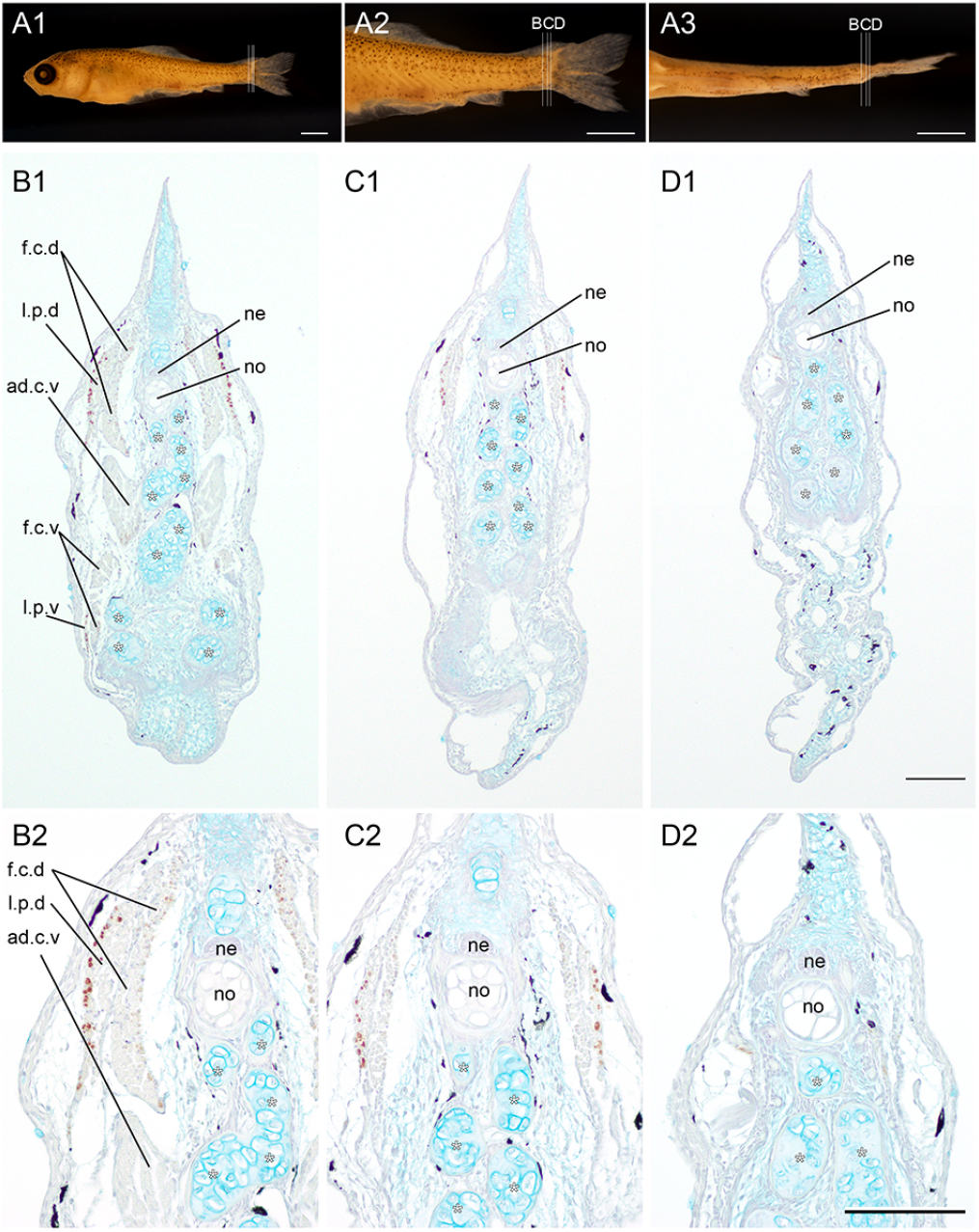
Distribution pattern of the slow muscle fibers in the *23 single #03* of the lab strain goldfish. (A). Whole body (A1), magnified lateral (A2), and ventral view (A3) of goldfish (#2023-0417-09-#03; 8.59 mm in the standard length, 22 dpf). The different levels of the sections at the caudal region were immunostained by the slow muscle fiber specific antibody (F59). (B, C, D). The sectioned levels are indicated in A2 and A3. Panels in the second and third rows show the low-magnification (B1, C1, D1) and the high-magnification (B2, C2, D2) images. White asterisks indicate the ventral caudal skeleton complex including pural, hypural, and hermal spines. Abbreviations: ad.c.v, adductor caudalis ventralis; f.c.d, flexor caudalis dorsalis; f.c.v, flexor caudals ventralis; l.p.d, lateralis profundus dorsalis; l.p.v, lateralis profundus ventralis; no, notochord; ne, neural tube. Scale bars = 1 mm (A1, A2, A3), 100 *µ*m (D1, D2). Histological sections in the same row have the same magnifications.

**Fig 10.**
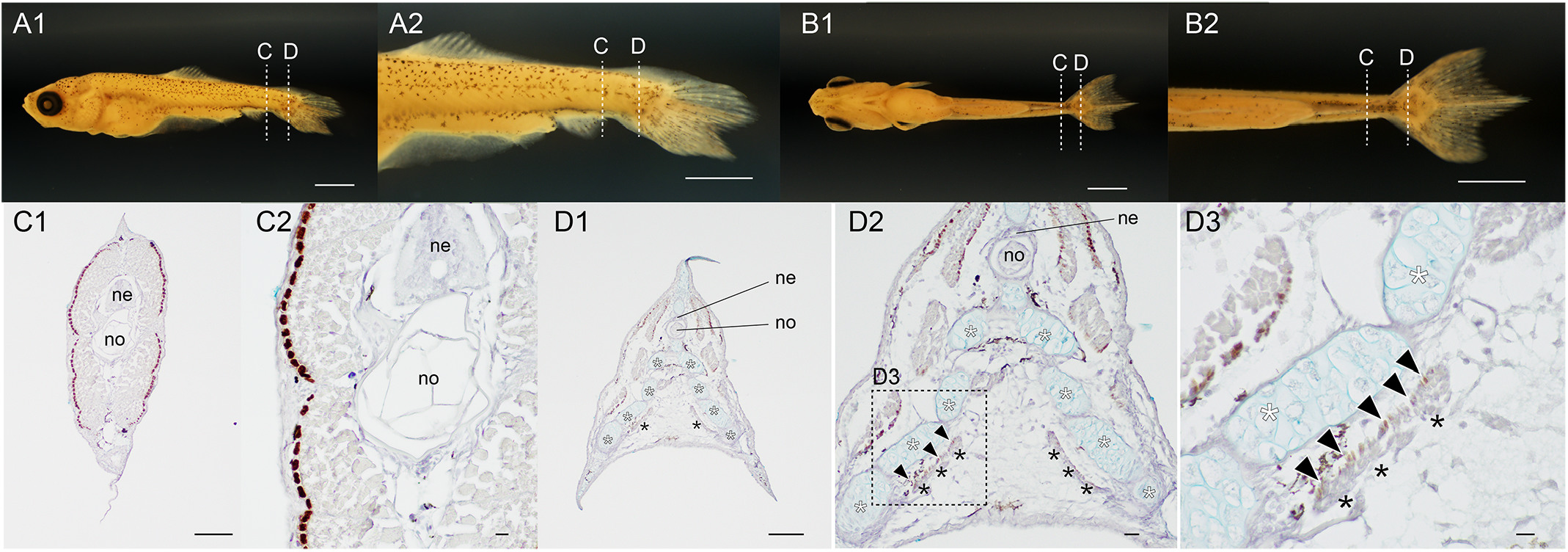
Distribution pattern of the slow muscle fibers in the *23 narrow twin #08* of the lab goldfish strain. (A, B). Whole body lateral (A1), magnified caudal lateral (A2), whole ventral (B1), and magnified caudal ventral views (B2) of goldfish (#2023-0417-09-#08; 7.16 mm, 22 dpf). (C, D). The different levels of the sections at the caudal region immunostained with the slow muscle specific antibody (F59). The sectioned levels are indicated in the panels of A and B. The first and second columns of section images are low (C1) and high (C2) magnified views at the trunk level. The third, second, and fifth columns of section images are low (D1), mid (D2), and high (D3) magnified views at the caudal level. The area of the high magnification view (D3) is outlined with the dashed box in panel D2. The muscular tissues located between the bifurcated caudal fin skeleton are indicated by black asterisks. White asterisks indicate the ventral caudal skeleton complex including pural, hypural, and hermal spines. Black arrowheads indicate slow muscle fibers in the flexor caudalis ventralis. Abbreviations: ne, neural tube; no, notochord. Scale bars = 1 mm (A, B). 100 *µ*m (C1, D1), 10 *µ*m (C1, C2, D2), 100 *µ*m (D3).

**Fig 11.**
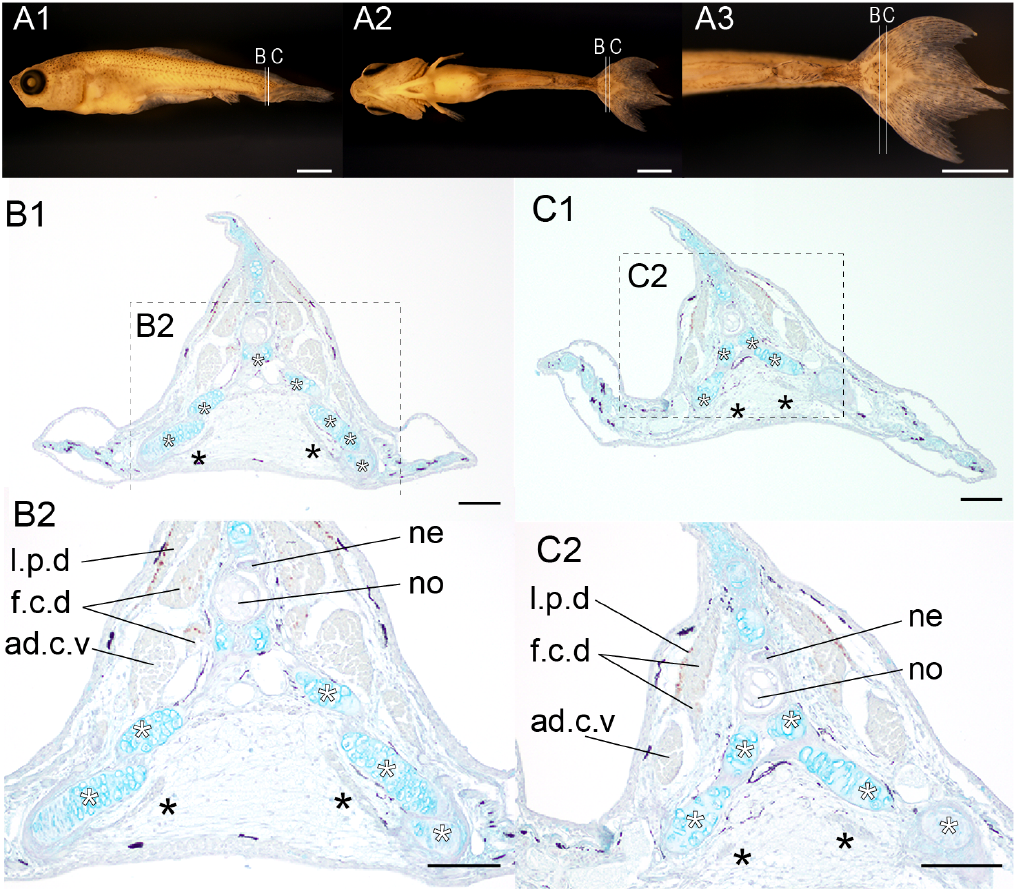
Distribution patterns of the slow muscle fibers in the *23 wide-twin #01* of the lab goldfish strain. (A). The lateral (A1), the ventral (A2), and the magnified ventral view (A3) of the goldfish (#2023-0417-09-#01, 22 dpf, 7.83 mm in the standard length). (B-C). The different levels of the sections at the caudal region immunostained with the slow muscle fiber specific antibody (F59). The sectioned levels are indicated in A1. The magnified views are indicated by the dashed boxes with panel labels. White asterisks indicate the ventral caudal skeleton complex including pural, hypural, and hermal spines. The muscular tissues located between the bifurcated caudal fin skeleton are indicated by the black asterisks in panels B1, B2, C1 and C2. Abbreviations: ad.c.v, adductor caudalis ventralis; f.c.d, flexor caudalis dorsalis; l.p.d, lateralis profundus dorsalis; ne, neural tube; no, notochord. Scale bars = 1mm (A1, A2, A3), 100 *µ*m (B1, B2, C1, C2)..

**Fig 12.**
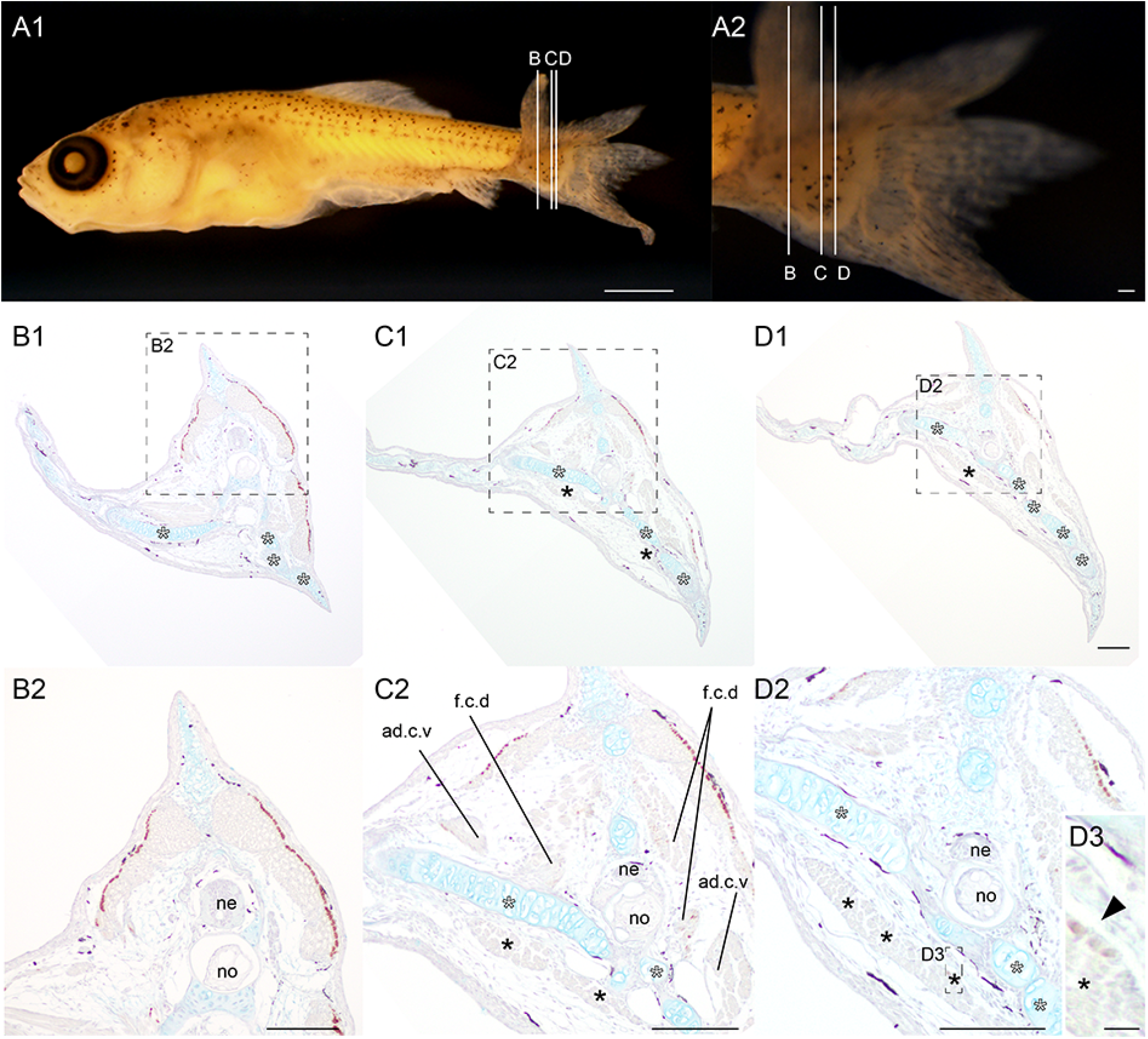
Distribution pattern of the slow muscle fibers at the *23 twisted #04* of the lab goldfish strain. (A). The whole body (A1) and the magnified view of the caudal region (A2) of the goldfish (#2023-0417-09-#04; 8.59 mm; at 22 dpf). (B-C). Different levels of the sections immunostained with the slow muscle fiber specific antibody (F59). The magnified views of the sections are indicated by dashed line boxes with panel labels. White asterisks indicate the ventral caudal skeleton complex including pural, hypural, and hemal spines. The muscular tissues located between the bifurcated caudal fin skeleton are indicated by black asterisks in panels C1, D1, C2, and D2. The black arrowhead in panel D3 indicates slow muscle fibers in the medial caudal muscle. Abbreviations: ad.c.v, adductor caudalis ventralis; f.c.d, flexor caudalis dorsals; ne, neural tube, no, notochord. Scale bars = 1 mm (A1), 100 *µ*m (A2, B2, C2, D1, D2). 100 *µ*m (D3). The panels in the second row are at the same magnification.

**Fig 13.**
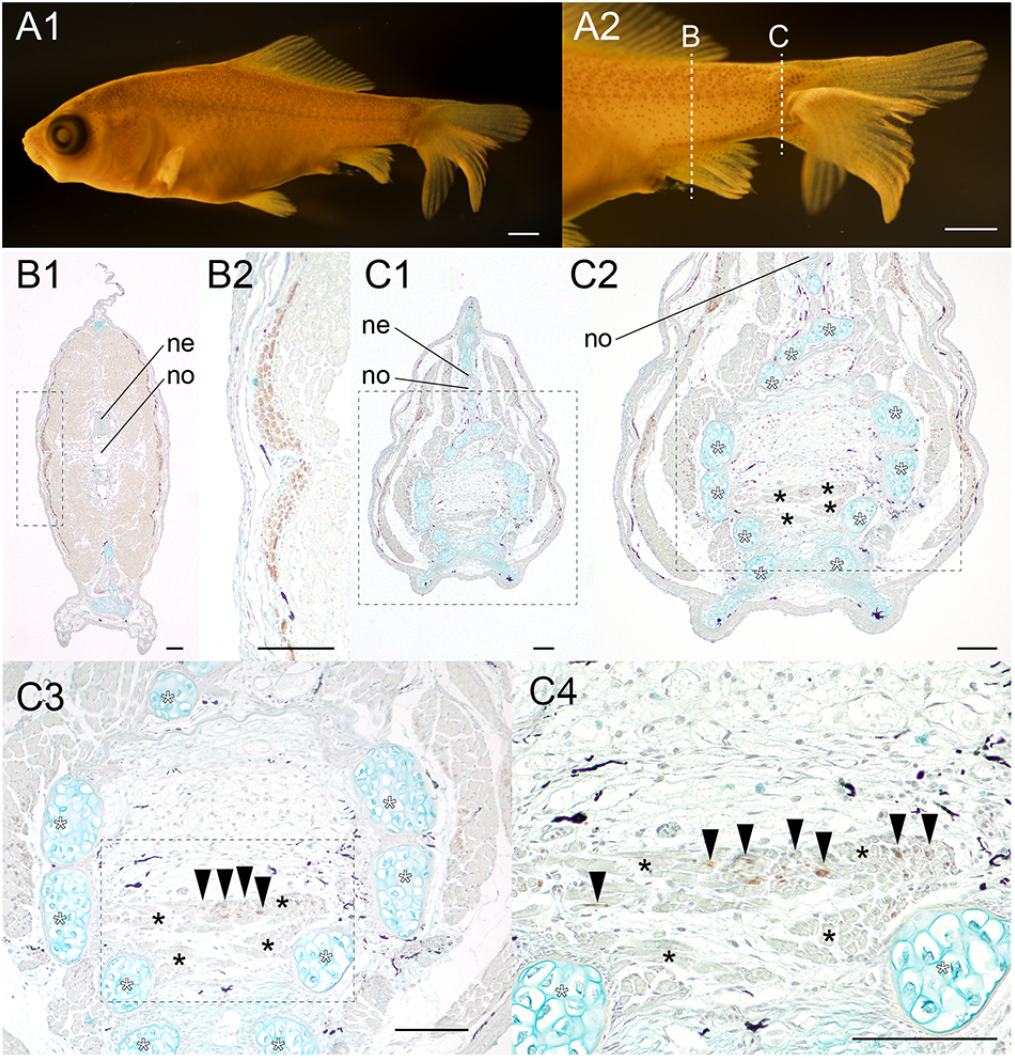
Distribution pattern of the slow muscle fibers at the *24 twin #01* of the lab goldfish strain. (A). The whole body (A1) and the magnified view of the caudal region (A2) of the goldfish (#2024-0419-03-#01; 13.07 mm; at 23 dpf). (B-C). Different levels of the sections immunostained with the slow muscle fiber specific antibody (F59). The magnified views of the sections are indicated by dashed line boxes with panel labels. White asterisks indicate the ventral caudal skeleton complex including pural, hypural, and hemal spines. The medial caudal muscle fibers are indicated by black asterisks in panels C2, C3, and C4. The black arrowhead in panel D3 indicates slow muscle fibers in the medial caudal muscle. Abbreviations: ne, neural tube, no, notochord. Scale bars = 1 mm (A1), 100 *µ*m (A2, B, C).

**Fig 14.**
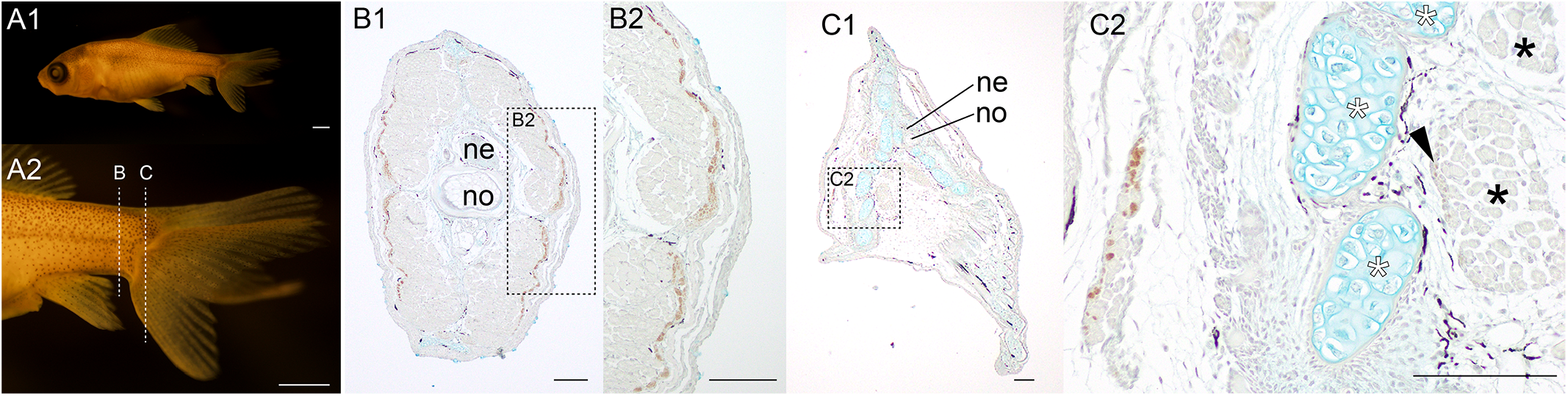
Distribution pattern of the slow muscle fibers in the *24 twin #02* of the lab goldfish strain. (A, B). Whole body lateral (A1), magnified caudal lateral (A2) of goldfish (#2024-0419-03-#02; 13.09 mm, 23 dpf). (B, C). The different levels of the sections at the caudal region immunostained with the slow muscle specific antibody (F59). The sectioned levels are indicated in the panels of A2. The area of the high magnification views (B2 and C2) are outlined with the dashed boxes in panel B1 and C1, respectively. The medial caudal muscle fibers are indicated by black asterisks in panel C2. White asterisks indicate the ventral caudal skeleton complex including pural, hypural, and hermal spines. Black arrowheads indicate slow muscle fibers in the flexor caudalis ventralis. Abbreviations: ne, neural tube; no, notochord. Scale bars = 1 mm (A, B). 100 *µ*m (C1, D1), 10 *µ*m (C1, C2, D2), 100 *µ*m (D3).

The following describes the morphological characteristics of these *chdS* lab strain individuals. The *23 single #03* did not have a clear bifurcated caudal fin, but several of its caudal skeletons were bifurcated (Fig. 9). The *23 narrow twin #08* had a bifurcated caudal fin (Fig. 10A1-10B2), although more than half of the area at the upper caudal fin was not bifurcated (Fig. 10A2). The *23 wide-twin #01* had a wellbilaterally bifurcated caudal fin, displaying a bifurcated upper fin lobe (Fig. 11A1-11A3). The *twisted #04* had a wellbifurcated but twisted caudal fin (Fig. 12A1-12A2), and a significant amount of muscular tissues was observed, differing from the *single #03* and the *narrow twin #08* (Fig. 12B-12D). The *24 twin #01* and *#02* showed the conventional bi- furcated caudal fin in their external morphology (Fig. 13A, 14A). The aforementioned morphological variations among these lab strain individuals facilitate the investigation of the relationship between their external caudal morphology and histological characteristics.

Since the slow muscle fibers at the surface muscle in all of these lab-strain were clearly detected, it is certain that the immunohistochemical analyses we performed provide the plausible results of the distribution patterns of the slow muscle fibers in the medial caudal muscles in these lab-strain individuals (Fig. 9B-D, 10C, 11B, 12B-D, 13B, 14B). Among these lab strains, the *23 narrow twin #08, 23 twisted #04, 24 twin #01*, and *24 twin #02* showed slow muscle fibers in their medial caudal muscles (the black arrows in Fig. 10C, 12B-D, 13B, 14B). Due to the ease of comparing the morphologies of the medial caudal muscles, it is clearer that the slow muscles fibers in the medial caudal muscles of the *23 narrow twin #08* were equivalent to those in ornamental goldfish (Fig. 6G4, 8B, 10D). On the other hand, although the comparison was more difficult in the *23 twisted #04* and the *24 twin #02* due to significant differences in muscle morphology, a common feature was found in that slow muscle fibers were distributed close to the caudal skeleton in these individuals (Fig. 12B-D, 14C).

Unlike the aforementioned twin-tail lab strain individuals, *24 twin #01* displays slow muscle fibers in the medial caudal muscles that are randomly scattered, rather than exhibiting a lateral bias (Fig. 13C2, C3, C4). The presence of slow muscle fibers is observed not only in the region near the bifurcated caudal skeletal elements but also in the medial part of the caudal skeleton. Due to the varied sectioning levels and distribution patterns of muscle fibers, resulting from the morphological diversity of these lab strain goldfish, the resolution of our histological comparison may be limited. For example, although we could not find the median caudal muscle in the 23 wide-twin 01, it is still uncertain whether this individual lacks the slow muscle fibers in the medial caudal muscle or not (Fig. 11). Nevertheless, it is clear that the distribution patterns of slow muscle fibers vary across these lab strains (black arrowheads in Fig. 10D3, 12D3, 13C4, 14C2).

## DISCUSSION

### Conserved slow muscle fiber distributions

Our study revealed the distribution patterns of the different types of muscle fibers at the caudal regions in the wild-type goldfish, zebrafish, and different types of twin-tail goldfish strains (including *Ryukin, Oranda*), and the lab strain *chdS* mutant goldfish. The successful implementation of immunohistochemistry in our study provides a unique opportunity to compare the distribution patterns of slow muscle fibers among these fishes.

We confirmed that the surface muscles at the mid-trunk region are covered by the slow muscle fibers in goldfish (Fig. 2-4, 6-14), similar to conventional teleost species (Kardong, 2006; Keenan and Currie, 2019; Liem et al., 2001). Additionally, our observations revealed the presence of slow muscle fibers along the proximal side of the deep muscles in the caudal region of the wild-type goldfish (Fig. 2-4, Fig. 15A). Given the similar distribution patterns of the slow muscle fibers observed in the zebrafish (Fig. 5 and 15A), it is reasonable to infer that this distribution pattern of slow muscle fibers within the deep muscles of the caudal region has been evolutionarily and developmentally conserved across these closely related teleost species. From the evidence that slow muscle fibers were found in a deep muscle in the caudal region of several teleost species from different lineages, it is expected that there may be a developmental mechanism specific to teleost fish that differentiates slow muscle fibers in the deep muscles at the caudal level (Flammang, 2014; Flammang and Lauder, 2008, 2009; Kryvi et al., 2021; Nag, 1972). Further comparison of immunohistochemical features of slow and fast muscle fibers in more distantly related teleost species will reveal whether this is correct. Moreover, although the caudal muscle in shark species has as well been shown to contain slow muscle fibers (Flammang, 2010), its distribution pattern is yet to be carefully compared, especially at the level of immunohistochemistry, with that of the slow muscle fibers in teleost species.

**Fig 15.**
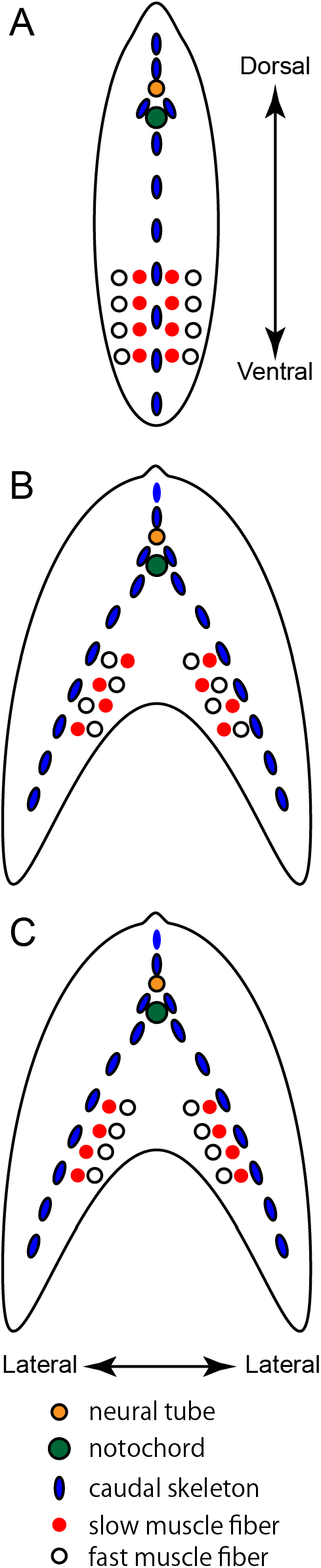
Schematic representation of the distribution patterns of muscle fibers. Transverse view at the caudal levels of wild-type goldfish and zebrafish (A), that of the mutant showing randomly mixed distribution patterns of the slow muscle fibers (see also Fig. 13), and that of conventional ornamental twin-tail goldfish (C). For simplicity, only the deep ventral muscles are shown.

We also identified the slow muscle fibers at the medial caudal muscles in *Ryukin, Oranda*, and four of investigated lab strain twin tail goldfish (Fig. 6G3-G4, 7B1-B3, 8B4- B5,10D3, 12D3, 13C4, 14C2). Based on the distributing area of the slow muscle fibers in the transverse plane, we could categorize their distribution pattern into two types; the randomly distributing type (Fig. 15B), and the biased distribution patterns (Fig. 15C). In our present research, the *24 twin #01* exhibits the randomly distributed slow muscle fibers in its medial caudal muscle (Fig. 13C4, 15B). On the other hand, *Ryukin, Oranda*, the *23 narrow twin #08, 23 twisted #04*, and *24 twin #02* can be categorized as the biased distribution pattern type (Fig. 6, 8, 10D3, 12D3, 14C2). Although there were differences in the morphology of the medial caudal muscle tissues, the slow muscle fibers in these individuals were distributed in close proximity to the caudal skeleton (Fig. 15C).

Among these individuals showing the biased distribution patterns of the slow muscle fibers in the medial caudal fin, *Ryukin, Oranda*, and the *23 narrow twin #08* enable us to consider the relationship between the medial caudal muscle and the other muscles. The observed slow muscle fibers in the twin-tail ornamental goldfish strain and the *23 narrow-twin #08* showed morphological resemblance with the slow muscle fibers in the flexor caudalis ventralis in wild-type goldfish (Fig. 2G2, 3C6, 4C2, 6G3, 6G4, 15). These resemblance of the distribution patterns of the slow muscle fibers in the twin-tail and wild-type goldfish suggest that a common underlying molecular mechanism and/or developmental process contribute to the formation of these muscles. Based on this interpretation, the randomly distributed slow muscle fibers in the *24 twin #01* led us to assume that the commonly underlying molecular developmental process might be disrupted, providing an opportunity to consider how the muscle tissues react to the morphological changes caused by the strong selective pressures (Fig. 13, 15C).

### Selective pressures and developmental mechanisms

The comparison of the ornamental twin-tail goldfish, the *narrow twin #08*, and *24 twin #01* offered a basis for considering a more plausible evolutionary scenario for the formation of the mediolaterally biased distribution patterns of slow muscle fibers in the medial caudal muscles (Fig. 6, 7, 10, 13 15). Two evolutionary scenarios have been considered to advance this discussion; whether the biased distribution patterns of slow muscles in the medial caudal fin are a consequence of i) selective pressures, or ii) conserved developmental mechanisms. Although the relationship between the former and the latter is not entirely mutually exclusive, this classification will contribute to advancing the discourse.

The former selection-based evolutionary scenario can be examined from previous comparative physiological and anatomical studies (Bernal et al., 2001; Blake et al., 2009; Fierstine and Walters, 1968; Shadwick et al., 2002). It was observed that twin-tail morphotype goldfish exhibit lower swimming performance compared to wild-type goldfish and this correlated with a reduced proportion of slow muscle fibers in twin-tail morphotype goldfish (Blake et al., 2009). Moreover, studies on the tuna fish group, known for highspeed cruising swimming, further demonstrated that there is a correlation between swimming performance and the position and quantity of slow muscle fibers (Bernal et al., 2001; Fierstine and Walters, 1968; Shadwick et al., 2002). From these findings, it is implied that the distribution patterns of the slow muscle fibers are highly evolvable and plastic traits. Thus, one may suggest the potential applicability of the selection-based evolutionary scenario as follows; if the mediolaterally biased distribution patterns of the slow muscle fibers are advantageous, the distribution patterns can be easily formed, even in a newly appeared muscle such as the medial caudal muscle. In essence, the specific distribution patterns of these two different muscle fibers seems to indicate adaptive significance. Under this selection-based evolutionary scenario, the random distribution patterns of the slow muscle fibers in the *24 twin #01* might be understood as the intermediate status (Fig 13, 15B).

However, applying the selection-based evolutionary scenario to the distribution patterns of slow muscle fibers in twin-tail goldfish is challenging due to the paucity of its supportive evidence in our study. In other words, we could not identify any clear advantageous points of the bilateral distribution patterns of slow muscle fibers in the medial caudal muscle of the twin-tail ornamental goldfish (Fig. 6G3, 6G4, 6B2, 7B3, 8B4, B5). Of course, one could argue that the presence of slow muscle fibers in the medial caudal muscles of the twin-tail ornamental goldfish contributes to maintaining both the open and closed states of the bifurcated caudal fin, thereby aiding in expressing a twin-tail phenotype preferred by breeders and fanciers. Indeed, if the distribution of slow muscle fibers is related to the spreading of the caudal fin, it is reasonable to consider that the current distribution patterns of the slow muscle fibers are a result of artificial selection. Nevertheless, it prompts skepticism regarding the applicability of this selection-based evolutionary scenario clarifying the laterally biased distribution patterns of slow muscle fibers, e. g. as observed in the *narrow twin #08* goldfish (Fig. 6G3, 6G4, 6B2, 6B3, 10D2, 10D3). The reasons for this skepticism are explained as follows.

Firstly, given that the *narrow twin #08* goldfish originated from the single-tail common goldfish strain, it seems improbable that these goldfish strains have undergone selective pressures leading to the development of twin-tail morphology with laterally biased distribution patterns of slow muscle fibers in the deep muscles of the caudal region. Secondary, the random distribution patterns of the slow-muscle fibers in the *24 twin #01* seems to be quite irregular distribution patterns, rather than the ancestral intermediate condition (Fig. 13C4). More specifically, the running direction of the muscle fibers of this individual are not well arranged in comparison with the other lab strains (Fig.6G4, 8B4, B5, 10D3, 13C4). In addition, the caudal skeleton of *24 twin #01* is positioned to surround the medial caudal muscle from both the dorsal and ventral sides, showing an exceptional and unique arrangement of the caudal skeleton (Fig. 13C2, C3).

Therefore, rather than relying solely on an adaptive explanation, it is pertinent to explore the possibility that the laterally biased distribution patterns of slow muscle fibers in the twin-tail ornamental goldfish and the *narrow twin #08* of the lab strain primarily reflect pre-existing molecular developmental mechanisms conserved in wild-type goldfish and zebrafish (Fig. 5B, 5C, 6G3, 6G4, 10D, 15BC). To put it differently, the lateral expansion of the caudal skeleton could have played a significant role in forming the laterally biased distribution patterns of slow muscle fibers in the medial caudal fin. When including the role of selection in this discussion, it seems more reasonable to consider that selection contributed to the elimination of irregular individuals, such as those with a caudal fin skeleton like the *24 twin #01*, rather than to the creation of a new molecular developmental mechanisms (Fig. 15BC).

### Notochord and muscle fiber distribution

From our present studies, we are unable to pinpoint the molecular developmental mechanisms responsible for the mediolaterally biased distribution patterns of the slow muscle fibers observed in the medial caudal muscle (Fig. 15C). However, by focusing on the function, position, and quantity of the notochord, we may approach the common underlying molecular developmental mechanisms that dictate the biased distribution of muscle fibers in both wild-type and twin-tail goldfish. Despite the presence of duplicated caudal musculoskeletal systems in twin-tail goldfish (including *Ryukin, Oranda*, and the lab strain, and previously investigated goldfish (see Li et al., 2019), our findings indicate that the notochord remains unduplicated (Fig. 1C, 6E1-6G5, 10D1) (see also Watase, 1887; Bateson 1894). These findings prompt an inquiry into the developmental origins of mediolaterally biased distribution patterns of slow muscle fibers within the bifurcated caudal muscular systems of twin-tail goldfish; how can a single notochord-derived signaling source induce similar distribution patterns of caudal muscle fibers in both single and twin-tail goldfish?

To address this inquiry, we need to consider two alternative hypotheses. The first suggests that the introduction of slow muscle fibers in the deep caudal region is governed by molecular mechanisms operating at the mid-trunk level, involving notochord-derived signals, as previously reported (Devoto et al., 1996; Du et al., 1997; Hadzhiev et al., 2007; Keenan and Currie, 2019; Siomava and Diogo, 2018; Stickney et al., 2000; Tanaka et al., 2023). The second proposes that the slow muscle fibers in the deep caudal region are defined by molecular mechanisms which differ from the aforementioned trunk notochord-derived signals (for example, see Elworthy et al., 2008). Although further studies are warranted, it is crucial to determine whether the former or latter hypothesis is more plausible, guiding future investigations.

For consideration of the first hypothesis, several points concerning the topological relationship between somite derivatives and the notochord must be addressed. Specifically, we cannot offer a plausible explanation for the observed mediolaterally biased distribution patterns in the flexor caudalis of the wild-type goldfish and the medial caudal muscle of twin-tail morphotype goldfish solely based on developmental mechanisms reliant on notochord-derived signals (Fig. 2G2, 3C5, 6G4, 10D3). While recognizing that somite rotation occurs during embryonic development (Hollway et al., 2007), one could propose an explanation for the varying distribution patterns of different muscle fiber types at the trunk and caudal levels. By altering the timing of somite rotation, different types of muscle tissues displaying varied distribution patterns of muscle fibers can be generated. However, it remains unclear how the same explanation can be applied to the histological features of the bifurcated caudal fin in twin-tail goldfish, given the uncertain developmental timing and origin of the medial caudal muscles (black arrowheads in Fig. 6G3, 10D). Thus, if we consider the mediolateral distribution patterns based on the already known molecular developmental mechanisms from the study of the trunk notochord, further examination of the developmental origin of the medial caudal muscles is required. Furthermore, the role of notochord-derived signals, including long-range and longduration retaining signals, in twin-tail goldfish should also be investigated (Hadzhiev et al., 2007; Tanaka et al., 2023).

In contrast to the first hypothesis centered on the molecular mechanisms found in the trunk notochord, the second hypothesis appears to offer a constructive and promising explanation for the laterally biased distribution patterns of slow muscle fibers in the medial caudal muscles. It is worth noting that slow muscle fiber differentiation can occur through various molecular mechanisms (Elworthy et al., 2008) and several caudal muscles develop differently from mid-trunk muscles (Siomava and Diogo, 2018). These studies suggest that the slow muscle fibers in both the flexor caudalis ventralis muscles of wild-type goldfish and the medial caudal muscles of twin-tail goldfish differentiate without the direct influence of notochord-derived signals. Given that the slow muscle fibers in the medial caudal muscle are adjacent to the caudal axial skeletons, it could be that axial skeleton-oriented signals contribute to the induction of the slow muscle fibers (the blue ellipses in Fig. 15). Assuming a relationship between the distribution patterns of slow muscle fibers and the location of the caudal skeleton, the randomly distributed slow muscle fibers in the *24 twin #01* might be explained as follows; the space enclosed by the caudal skeleton could be filled with signals in an unusual manner, leading to a lack of specific polarity and resulting in the random appearance of slow muscle fibers (Fig 13C).

Moreover, other signal sources might influence the differentiation of the slow and fast muscle fibers in goldfish. During embryonic development, the embryonic tailbud harbors different types of molecular signals, and its cells differentiate into multiple derivatives, including the precursors of muscle tissues (Agathon et al., 2003; Das et al., 2019; Henrique et al., 2015; Lawton et al., 2013; Ohta et al., 2007; Row et al., 2016). Notably, at least some known signaling domains are bifurcated in the tailbud of twin-tail goldfish, despite it having a single notochord (see *bmp4* in Abe et al., 2014, a regulator of muscle specification in zebrafish, Esterberg et al., 2008). It also very well could be that the function of the notochord and its attached tissues at the most posterior level differs from that of equivalent tissues (notochord and neural tube) at the mid-trunk region. Indeed, it has been shown in salmon species (*Salmo salar*) that the posterior- and anterior regions of the notochord could be very different (Kryvi et al., 2021). Thus, it is reasonable to consider that applying the same molecular mechanisms directly to explain the differentiation processes of both trunk and tail muscle fibers could be challenging. Further investigations into the developmental origin of different types of muscle fibers in the tail region will help elucidate whether the first or second hypothesis is more tenable.

### Considerations on novel morphology

Our observations also shed light on the relationship between the *chordin* gene mutation and the distribution pattern of slow muscle fibers in goldfish. Mediolaterally biased distribution patterns of slow muscle fibers were observed in the deep muscles of different types of goldfish (Fig. 2D1, 3B-C6, 6DG, 7B, 8, 10D2). Since these goldfish individuals are varied in the *chordin* gene genotype, it is reasonable to expect that the absence of the *chdS*^*wt*^ allele does not significantly alter the topological relationship between slow and fast muscle fibers in most of examined goldfish individuals. It also indicates that caudal muscle developmental mechanisms can generate similar types of muscular tissues even under modified dorsal-ventral patterning. Thus, the appearance of similar distribution patterns of slow muscles results from developmental systems independent of the *chdS* gene. Consequently, the conserved distribution patterns of slow muscle fibers in different types of fishes (including wild-type goldfish, zebrafish, *Ryukin, Oranda*, and the lab strain goldfish with the *chdS*^*E127X/E127X*^ genotype) stem from partially integrated or parcellated developmental mechanisms that can produce novel morphologies with quasi-identical histological characteristics.

Based on the results of this study, the existence of molecular mechanisms that generate characteristic muscle tissue features in twin-tail goldfish, independent of the *chdS* gene, is suggested. These features not only resemble external structures but also homologous histological structures at the tissue level between wild-type and twin-tail goldfish at the caudal region. While our focus in this study centered on goldfish caudal fin histology, our findings provide insightful perspectives for contemplating the emergence of largescale morphological evolution. For example, the appearance of paired fins and paired nostrils are quite similar to the appearance of the bifurcated caudal fin in the twin-tail goldfish (Abe et al., 2007; Abe and Ota, 2016; Gai et al., 2022; Janvier and Arsenault, 2007; Kuratani, 2012; Kuratani et al., 2016; Tzung et al., 2023). When these left-right bilaterally duplicated homologous morphologies emerge, they simultaneously give rise to novel medial morphology between the left and right caudal skeletons. In the process of their appearance, pre-existed molecular developmental systems are employed with modifications for the formation of novel tissues in the medial morphologies. Our current goldfish study provides an experimental case aimed at understanding how tissues and cells behave in the newly emerged space that arises when novel morphological features evolve.

## CONCLUSIONS

The present work shows that the deep muscles at the caudal level in wild-type goldfish and zebrafish exhibit the laterally biased distribution patterns of the slow muscle fibers, indicating that these distribution patterns might be derived from long-term conserved molecular developmental mechanisms. Similar distribution patterns of the slow muscle fibers were observed at the medial caudal muscles in the bifurcated caudal fin of the twin-tail goldfish. Their similarity implies that the distribution patterns of the slow muscle fibers in these different muscles are organized by the same molecular developmental mechanisms. Our present study provides an empirical example to consider how histology-level phenotypes are influenced by selective pressure and conserved developmental mechanisms.

## ACKNOWLEDGEMENTS

We are grateful to Wen-Hui Su (SHUEN-SHIN Breeding Farm) and You Syu Huang (ex-member of the Aquaculture Breeding Institute, Hualian), for technical advice on goldfish breeding in Taiwan. We also thank the following current and ex-members of the Yilan Marine Research Station, Institute of Cellular and Organismic Biology; the late Hung-Tsai Lee, Chia-Chun Lee, Chihi-Chiang Lee, and Tsai Han Chuan for maintenance of aquarium systems; Chi-Fu Hung, Jhih-Hao Wei, and Fei Chu Chen for administrative support; Teng Yu-Feng (Big Wind Technology Co. LTD) and Shi-Chieh Liu for designing aquarium systems. We also thank the Taiwan Zebrafish Core Facility at Academia Sinica.

## AUTHOR CONTRIBUTIONS

**Conceptualization**: Kinya G. Ota 太田欽也 (KGO). **Funding Acquisition**: KGO. **Investigation**: KGO, Tzu-Chin Chi 紀子勤 (TCC). **Methodology**: KGO, TCC, Chen-Yi Wang 王貞懿(CYW), Ing-Jia Li 李穎佳(IJL). **Project Administration**: KGO. **Resources**: CYW, IJL, KGO. **Supervision**: KGO. **Image acquisition**: KGO. **Writing – Original Draft**: KGO. **Writing – Review & Editing**: KGO, Paul Gerald Layague Sanchez (PGLS), IJL, TCC.

